# Deletion of *Nudt19* Increases Albuminuria in Mice Fed a High Fat Diet

**DOI:** 10.1101/2025.03.22.644727

**Authors:** Dominique C. Saporito, Rachel D. King, Schuyler D. Vickers, Emily A. Wyda, Sruthi Balaji, Judy A. King, Roberta Leonardi

## Abstract

Nudix hydrolase 19 (NUDT19) is a peroxisomal enzyme that hydrolyzes CoA species at the phosphodiester bond and has been linked to peroxisomal dysfunction in the context of diabetic kidney disease. Despite its predominant expression in mouse kidneys, the physiological role of NUDT19 remains poorly understood. To investigate its function under metabolic stress, we fed *Nudt19*^*-/-*^ mice a high fat diet (HFD) for 15 weeks. *Nudt19* deletion exacerbated HFD-induced albuminuria, suggesting a previously unrecognized role in kidney function. This phenotype was associated with altered lipid metabolism in the kidneys, including reduced levels of non-esterified fatty acids and specific mono-acyl lipids, as well as differential expression of proteins involved in lipid metabolism. These included ECH1, THIKB, and ECHD2, enzymes involved in peroxisomal and mitochondrial β-oxidation; C19orf12, a lipid droplet-associated protein; and the lipolysis-stimulated lipoprotein receptor (LSR). These findings highlight NUDT19 as a key regulator of renal lipid homeostasis and suggest that its loss contributes to kidney dysfunction under conditions of dietary lipid overload.

## Introduction

The kidneys are highly energy-demanding organs, consuming a substantial amount of ATP to sustain, among other functions, the tubular reabsorption of filtered solutes. This energy demand is met through the use of a variety of metabolic fuels, including glucose, fatty acids, and amino acids, with utilization patterns varying across different regions of the kidney and in response to physiological conditions [1-3]. With the exception of the inner medulla, where glucose is preferentially converted to lactate [4, 5], energy generation in the kidneys is largely provided by the mitochondria, which are highly enriched in proximal tubule (PT) cells [6, 7]. PT cells also contain a high density of peroxisomes [8, 9], organelles that plays a key role in lipid metabolism, including the synthesis of plasmalogens and the oxidation of branched-chain, dicarboxylic, and very long-chain fatty acids [10-15]. While peroxisomal fatty acid oxidation does not directly generate ATP, the unique ability of peroxisomes to process fatty acids that are not substrates for mitochondrial β-oxidation supports overall energy production by breaking them down into shorter products, which are then transferred to and completely oxidized in the mitochondria.

A large number of reactions involved in fuel metabolism require coenzyme A (CoASH), an essential cofactor serving as the main acyl carrier in mammalian cells. CoASH and its acyl-CoA thioesters are involved in the metabolism of carbohydrates, fatty acids, amino acids and ketone bodies, and the post-translational modifications of histones and other proteins [16-20]. Separate pools of this cofactor are found in the cytosol, mitochondria, and peroxisomes, where they support distinct metabolic pathways, as well as in the endoplasmic reticulum (ER) lumen and nucleus, where they contribute to processes such as protein quality control, autophagy, and gene regulation [21-23]. Within mitochondria, peroxisomes, and the ER, specific enzymes hydrolyze CoA species to 3’,5’-ADP and (acyl-)phosphopantetheine. This hydrolysis potentially regulates the size and composition of local CoASH/acyl-CoA pools, which in turn affects organelle-specific processes. For instance, liver-specific deletion of *Fitm2*, which encodes FIT2, an ER membrane protein that preferentially hydrolyzes long, unsaturated acyl-CoAs, disrupts ER homeostasis, leading to ER stress and liver injury [24, 25]. Similarly, alterations in the activity of NUDT7, a peroxisomal CoA-degrading enzyme highly expressed in the liver, affect peroxisomal processes. These include bile acid synthesis, whose final steps occur in these organelles, and the metabolism of dicarboxylic fatty acids [26, 27]. NUDT19 is another peroxisomal Nudix hydrolase with specific CoA-diphosphohydrolase activity [28, 29]. While NUDT19 and NUDT7 are both Nudix hydrolases, these enzymes share limited sequence similarity (∼30%) and exhibit distinct tissue distributions and regulatory properties. NUDT19 is almost exclusively expressed in mouse kidneys, where it was initially identified (originally named RP2p) as a protein whose transcript levels were strongly upregulated by androgens [30]. Unlike NUDT7, NUDT19 protein levels do not change between the fed and fasted states [29]. NUDT19 exhibits tubular staining in normal kidneys and, consistent with its subcellular localization, has been linked to peroxisomal dysfunction in diabetic kidney disease (DKD), with its reduction observed in both diabetic kidney mouse models and human DKD samples [31]. NUDT19 also contains a unique 45–49-amino acid insertion of unknown function within its Nudix box [28, 32], which may contribute to its distinct substrate specificity compared to NUDT7 [29]. These properties suggest that, while both are peroxisomal enzymes, NUDT7 and NUDT19 likely fulfill distinct organ-specific functions.

Deletion of *Nudt19* in mice fed regular chow does not elicit any overt phenotype [29], but the physiological role of NUDT19, particularly under metabolic stress, remains poorly defined. In this study, we used a high fat diet (HFD) to impose a fatty acid-mediated metabolic challenge on *Nudt19*^*-/-*^ mice. Under these conditions, deletion of *Nudt19* was associated with changes in lipid metabolism and exacerbated the albuminuria caused by HFD-feeding.

### Experimental Procedures

Reagents were purchased from the following suppliers: Etomoxir from Tocris Bioscience, ^14^C-palmitic acid from American Radiolabeled Chemicals, horseradish peroxidase-conjugated goat anti-rabbit secondary antibody from Thermo Fisher Scientific, MS-grade acetonitrile from Honeywell. All other chemicals were of analytical grade or better and were purchased from Sigma-Aldrich or Fisher Scientific, unless stated otherwise. The NUDT7, NUDT8 and NUDT19 antibodies were generated and validated as described previously [29, 33].

#### Animal studies

Embryos of *Nudt19tm1(KOMP)Vlcg* mice were purchased from the Knock-out Mouse Project (KOMP) repository and bred to generate *Nudt19*^*-/-*^ mice as previously described [29]. These mice were subsequently bred to β-actin-cre transgenic mice (B6N.FVB-*Tmem163*^*Tg(ACTB-cre)2Mrt*^/CjDswJ, Jackson Laboratory stock number 019099) to excise the floxed neomycin cassette and establish the line of *Nudt19*^*-/-*^ and control mice characterized herein. Genotype was assigned by multiplex PCR analysis using the Accustart II PCR Genotyping Kit (QuantaBio) and primers reg-Nudt19-R, reg-Nudt19-wtR and reg-Nudt19-wtF2 [29], in combination with primer reg-LacF (ACTTGCTTTAAAAAACCTCCCACA). PCR products of 195 and 685 base pairs indicated the presence of wild type or knockout alleles, respectively.

Mice were fed a standard chow diet (Tekland 2018S) and housed in an animal facility kept at a room temperature of 22.0 ± 0.2 °C, room humidity of 40% ± 2%, and a 12-h light/12-h dark cycle, with the dark cycle starting at 18:00 h. Starting at 6 weeks of age, wild type and *Nudt19*^*-/-*^ mice were randomly assigned to eat either a low fat control diet (CD, Research Diets D12450J, 10% kcal from fat) or a HFD (Research Diets D12492, 60% of calories from fat) for 15 weeks. Body weight monitoring was conducted every two weeks. Blood pressure measurements were obtained with a MRBP tail cuff blood pressure multi-channel system (IITC Life Science). The reported blood pressure measurements were obtained at week 12, after the mice were acclimated to the instrument and measurement procedure for one week. Body composition was analyzed with an EchoMRI™ instrument at week 13. Blood glucose was measured at week 14 either in the fed state or after fasting the mice for 24 h. Urine was collected from mice singly housed in metabolic cages during the last 24 h of a 6-day acclimation period started at week 14. For tissue harvest, the mice were anesthetized by administration of isoflurane, followed by blood collection by cardiac puncture and subsequent removal of the organs of interest. Tissue samples were quickly weighed before flash-freezing in liquid nitrogen, then stored at -80 °C until analysis. Blood samples were allowed to clot, before being centrifuged at 10,000 x g for 10 min to isolate serum. All studies were approved by the Institutional Animal Care and Use Committees of West Virginia University

#### Histology, lipids, and urine analyses

Kidneys were harvested from mice in the fed state at 9:00 AM, cut longitudinally in half, with one half immediately fixed in 10% neutral buffered formalin for 1 week at 4 °C. Fixed samples were embedded in paraffin blocks, cut into 5 μm sections, and stained with hematoxylin and eosin (H&E) by the WVU Histopathology Core Facility. Serum triglycerides and total cholesterol levels were measured using Stanbio kits (EKF Diagnostics USA), following the manufacturer’s instructions. Lipids were extracted from flash-frozen kidney samples as previously described [27], and re-suspended in 100 µl of 5% NP-40. This solution was then used to measure kidney triglycerides and total cholesterol levels, following the same protocol used for the serum samples. Urinary sodium was quantified with a MyBioSource colorimetric kit, as per manufacturer’s instructions. Thiobarbituric acid reactive species (TBARS) were measured in 40 µl of urine mixed with an equal volume of an 8% sodium dodecyl sulfate solution, as described by Ohkawa H., et al. [34].

#### Western blots

For Western blot analysis, flash-frozen kidney cortex samples were homogenized in ice-cold radioimmunoprecipitation assay buffer containing 1X protease inhibitors (Biotool), and centrifuged at 10,000 x g for 10 min at 4°C. Proteins (100 μg) were fractionated on 4–12% bis-Tris polyacrylamide gels, transferred onto PVDF membranes, and visualized by Ponceau S staining. The NUDT8 and NUDT19 antibodies were used at 1:3000 and 1:5000 dilution, respectively, and incubated at 4 °C overnight. The NUDT7 antibody was used at 1:3000 dilution and incubated at room temperature for 1 h. Horseradish peroxidase-conjugated goat anti-rabbit IgG was used as the secondary antibody at a 1:22,500 dilution, incubating for 1 h at room temperature. Antibody signals were detected by chemiluminescence on a G:BOX Chemi XX9 imaging system (Syngene). Densitometric analysis was performed utilizing ImageJ (https://imagej.nih.gov/ij/). The antibody signal in each sample was normalized to the signal corresponding to the total protein loaded, as determined by the intensity of the Ponceau S stain.

#### Total CoA and acyl-CoA analysis

Total CoA levels in kidney cortex samples were determined by hydrolyzing all acyl-CoAs to CoASH under alkaline conditions. This was followed by derivatization of CoASH with monobromobimane, HPLC separation, and detection with a fluorescence detector, as previously described [35]. Acyl-CoA analysis was conducted as previously reported [26].

#### Untargeted metabolomics and proteomics

Global metabolomic profiling, with associated statistical analyses and pathway enrichment assessments, were conducted by Metabolon, Inc. Peak intensities normalized to tissue weights and significantly changed metabolites are reported in **supplementary Table S1**.

Proteomics analysis was conducted by the IDeA National Resource for Quantitative Proteomics. Briefly, protein samples underwent reduction, alkylation, and chloroform/methanol extraction before digestion with trypsin. Peptides were labeled using a tandem mass tag 10-plex isobaric label reagent set and combined into one group, then fractionated into 46 fractions using a 100 x 1.0 mm Acquity BEH C18 column (Waters). These fractions were consolidated into 18 super-fractions and further separated by nano-LC on an XSelect CSH C18 2.5 um resin (Waters) on an in-line 150 x 0.075 mm column. Eluted peptides were ionized by electrospray followed by mass spectrometric analysis on an Orbitrap Eclipse Tribrid mass spectrometer using multi-notch MS3 parameters. MS data were acquired using the FTMS analyzer in top-speed profile mode at a resolution of 120,000 over a range of 375 to 1500 m/z. Following CID activation with normalized collision energy of 31.0 eV, MS/MS data were acquired using the ion trap analyzer in centroid mode and normal mass range. Using synchronous precursor selection, up to 10 MS/MS precursors were selected for HCD activation with normalized collision energy of 55.0 eV, followed by acquisition of MS3 reporter ion data using the FTMS analyzer in profile mode at a resolution of 50,000 over a range of 100-500 m/z. Proteins were identified and MS3 reporter ions quantified using MaxQuant (Max Planck Institute) against the UniprotKB *Mus musculus* database with a parent ion tolerance of 3 ppm, a fragment ion tolerance of 0.5 Da, and a reporter ion tolerance of 0.003 Da. Scaffold Q+S (Proteome Software) was used to verify MS/MS based peptide and protein identifications. Protein identifications were accepted if they could be established with less than 1.0% false discovery and contained at least 2 identified peptides. Protein probabilities were assigned by the Protein Prophet algorithm [36] to perform reporter ion-based statistical analysis. Protein TMT MS3 reporter ion intensity values were assessed for quality and normalized using ProteiNorm [37]. The data was normalized using VSN [38] and analyzed using proteoDA to perform statistical analysis using Linear Models for Microarray Data (limma) with empirical Bayes (eBayes) smoothing to the standard errors [39]. Proteins with an FDR adjusted p-value < 0.05 were considered to be significant. Ion intensities and significantly changed proteins are reported in **supplementary Table S2**.

#### Measurement of fatty acid oxidation

To assess fatty acid oxidation in kidney cortex slices, both kidneys from a single mouse were collected, the cortex was dissected, and finely minced using a razor blade. The kidney cortex slices were then portioned into approximately 20 mg aliquots and placed into round bottom 14 ml tubes containing 1 ml of ice-cold M199 medium containing carnitine at a final concentration of 450 µM and either etomoxir at a final concentration of 67.5 µM, or an equal volume of vehicle (DMSO). After pre-incubating the kidney slices for 25 minutes at 37°C with shaking, the reaction was initiated by adding a pre-warmed substrate mix composed of a combination of unlabeled and ^14^C-labeled palmitic acid complexed with bovine serum albumin (BSA) (at a ratio of 5:1, fatty acid:BSA). The final concentration of fatty acid was 100 µM (2 mCi/mmol specific activity) in the final volume of 1.5 ml. Duplicate reactions were incubated for 1 h or stopped immediately to assess background radioactivity. Reactions were halted by transferring 500 µl of the solution above the kidney slices to 1.5 ml tubes containing 50 µl of 70% perchloric acid. After a 1 h incubation at room temperature, the tubes were centrifuged at 13,000 x g for 10 minutes. A 300 µl aliquot of the resulting supernatants was transferred to scintillation vials, and the radioactivity associated with the acid-soluble metabolites (ASM) was quantified by scintillation counting in a PerkinElmer Tri-Carb 4810TR liquid scintillation analyzer.

## Results

### Effect of HFD-feeding on Nudt19^-/-^ mice

We previously reported that chow-fed *Nudt19*^*-/-*^ mice had no overt phenotype [29]. Given the localization of NUDT19 in kidney peroxisomes and the role of these organelles in lipid metabolism, we tested the effect of feeding a HFD on *Nudt19*^*-/-*^ mice and wild type controls for 15 weeks. As expected, mice fed the HFD gained significantly more weight and accumulated more fat mass than mice fed the CD, with no differences between genotypes (**Fig. 1**). HFD feeding also resulted in similar increases in fed and fasted blood glucose levels and serum cholesterol levels in both wild type and *Nudt19*^*-/-*^ mice (**Table 1**).

**Table 1.** Selected features of *Nudt19*^*-/-*^ (KO) and wild type (WT) mice fed the CD and HFD. Values are expressed as the mean of 5-23 mice per condition ± SEM. For measurements in the fasted state, food was withdrawn for 24 h. All other measurements were conducted on randomly fed mice. For urine analyses, mice were housed in metabolic cages for at least 6 days before collecting urine samples over 24 h. The # symbol indicates significant diet-induced differences within each genotype: ## *p* < 0.01, and ### *p* < 0.001. Two-way ANOVA. BW, body weight; TAG, triglycerides.

| Parameter | Genotype | Diet |  |
| --- | --- | --- | --- |
|  |  | CD | HFD |
| Blood glucose, mg/dl | WT | 93 ± 2 | 129 ± 5### |
| | KO | 90 ± 2 | 125 ± 7### |
| Blood glucose, fasted, mg/dl | WT | 57 ± 1 | 101 ± 4### |
| | KO | 56 ± 3 | 101 ± 6### |
| Blood nitrogen urea, mg/dl | WT | 27 ± 1 | 27 ± 2 |
| | KO | 23 ± 2 | 29 ± 3 |
| Serum TAG, mg/dl | WT | 66 ± 12 | 61 ± 3 |
| | KO | 67 ± 19 | 78 ± 5 |
| Serum cholesterol, mg/dl | WT | 126 ± 9 | 166 ± 8## |
| | KO | 103 ± 4 | 170 ± 7### |
| Liver weight, g | WT | 1.58 ± 0.06 | 2.39 ± 0.13### |
| | KO | 1.44 ± 0.06 | 2.33 ± 0.12### |
| Right kidney weight, mg | WT | 146 ± 6 | 149 ± 10 |
| | KO | 127 ± 3 | 147 ± 8 |
| Left kidney weight, mg | WT | 132 ± 5 | 159 ± 10# |
| | KO | 124 ± 3 | 141 ± 7 |
| Urine output, ml/24h | WT | 1.4 ± 0.2 | 1.3 ± 0.1 |
| | KO | 1.7 ± 0.2 | 1.1 ± 0.1 |
| Urinary TBARS, μM | WT | 11 ± 2 | 18 ± 2 |
| | KO | 12 ± 3 | 19 ± 1 |
| Urinary sodium, mM | WT | 82 ± 10 | 85 ± 5 |
| | KO | 88 ± 11 | 92 ± 8 |

**Figure 1.**
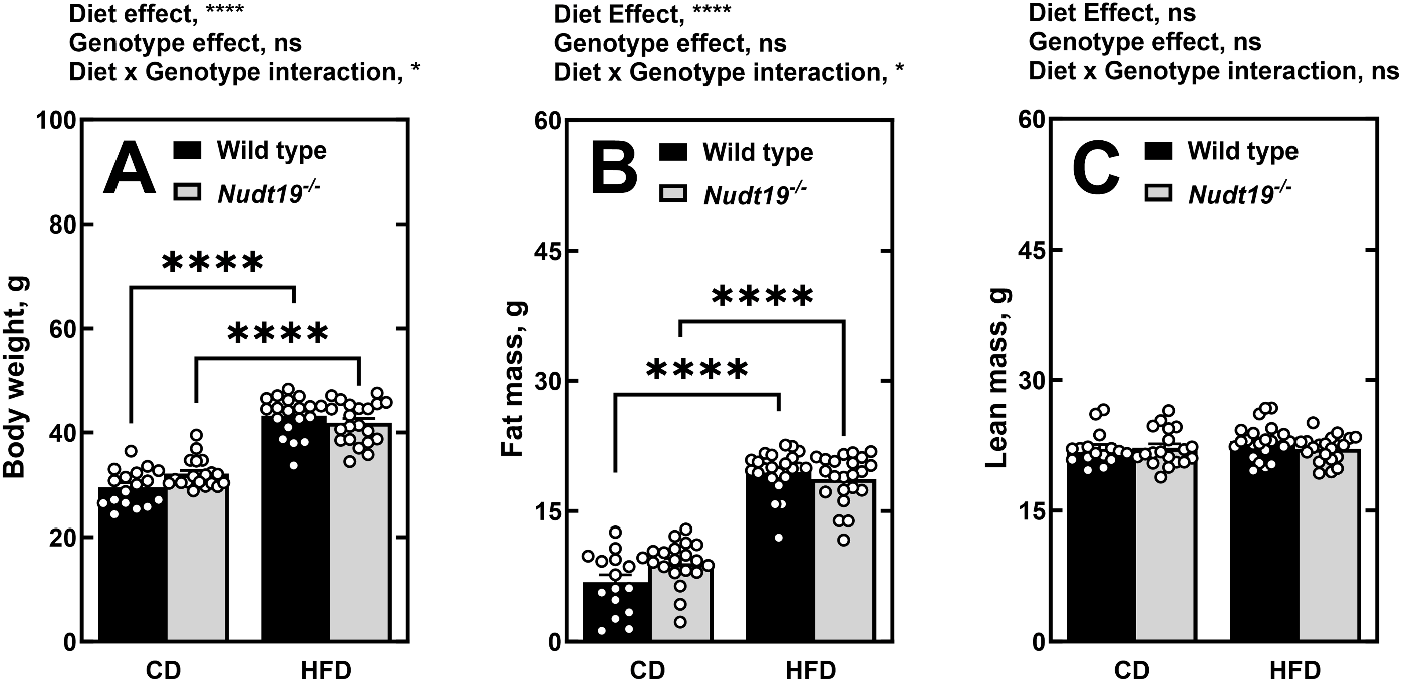
Effect of HFD on body weight and body composition. (A) Body weight, (B) fat mass and (C) lean mass of *Nudt19*^*-/-*^ and wild type mice after 13 weeks of CD and HFD feeding. Data are shown as the mean (bars) of measurements from individual mice (circles) ± SEM. Two-way ANOVA; ns, not significant, *, p<0.05; ****, p<0.0001.

In response to HFD feeding, wild type mice showed a significant decrease in NUDT19 and NUDT7 protein levels in the kidneys (**Supplementary Figure 1**). Regardless of diet, deletion of *Nudt19* did not cause a compensatory increase in NUDT7 or NUDT8, the two other Nudix hydrolases that degrade CoA species. Histological analysis of renal cortex areas revealed that both wild type and *Nudt19*^*-/-*^ mice fed the CD exhibited rare vacuolated tubular epithelial cells (1/4 of wild type mice and 0/7 of *Nudt19*^*-/-*^ mice analyzed) (**Fig. 2A**). After 15 weeks on the HFD, the frequency of vacuolated tubular epithelial cells increased in both wild type (6/6 mice) and *Nudt19*^*-/-*^ mice (6/6 mice), although triglycerides and total cholesterol levels remained overall unchanged by diet or genotype (**Fig. 2B-C**).

**Figure 2.**
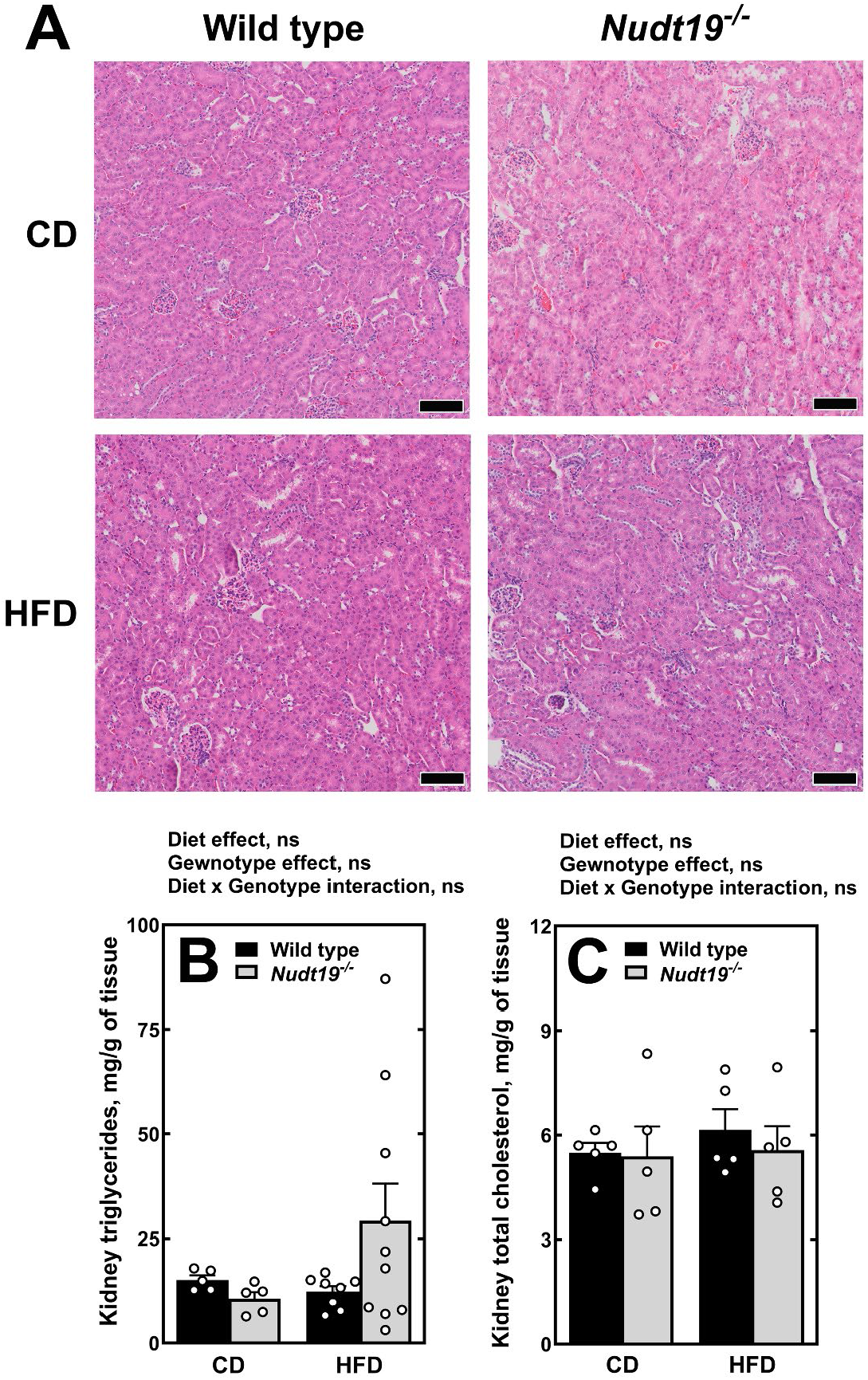
Histological analysis and kidney lipids. (A) Representative bright field images of H&E-stained kidney sections from wild type and *Nudt19*^*-/-*^ mice fed the CD and HFD. Bar = 100 µm. (B) Kidney triglycerides and (C) total cholesterol. Data are shown as the mean (bars) of measurements from individual mice (circles) ± SEM. Two-way ANOVA; ns= not significant.

To examine the effect of *Nudt19* deletion on kidney function we analyzed urine collected over 24 h for sodium, TBARS, a marker of lipid peroxidation, and albumin-to-creatine ratio (ACR). Sodium secretion was unaffected by diet and similar between genotypes (**Table 1**). TBARS tended to be higher in the urine of HFD-fed mice, regardless of genotype, but the difference was not statistically significant (**Table 1**). The concentration of albumin in the urine and the ACR significantly increased in mice fed the HFD and were found to be even higher in mice lacking NUDT19 (**Fig. 3**). Overall, these data indicated that deletion of *Nudt19* specifically exacerbated the deleterious effect of a HFD on albumin reabsorption in the kidneys. We then focused all subsequent analyses on mice fed the HFD.

**Figure 3.**
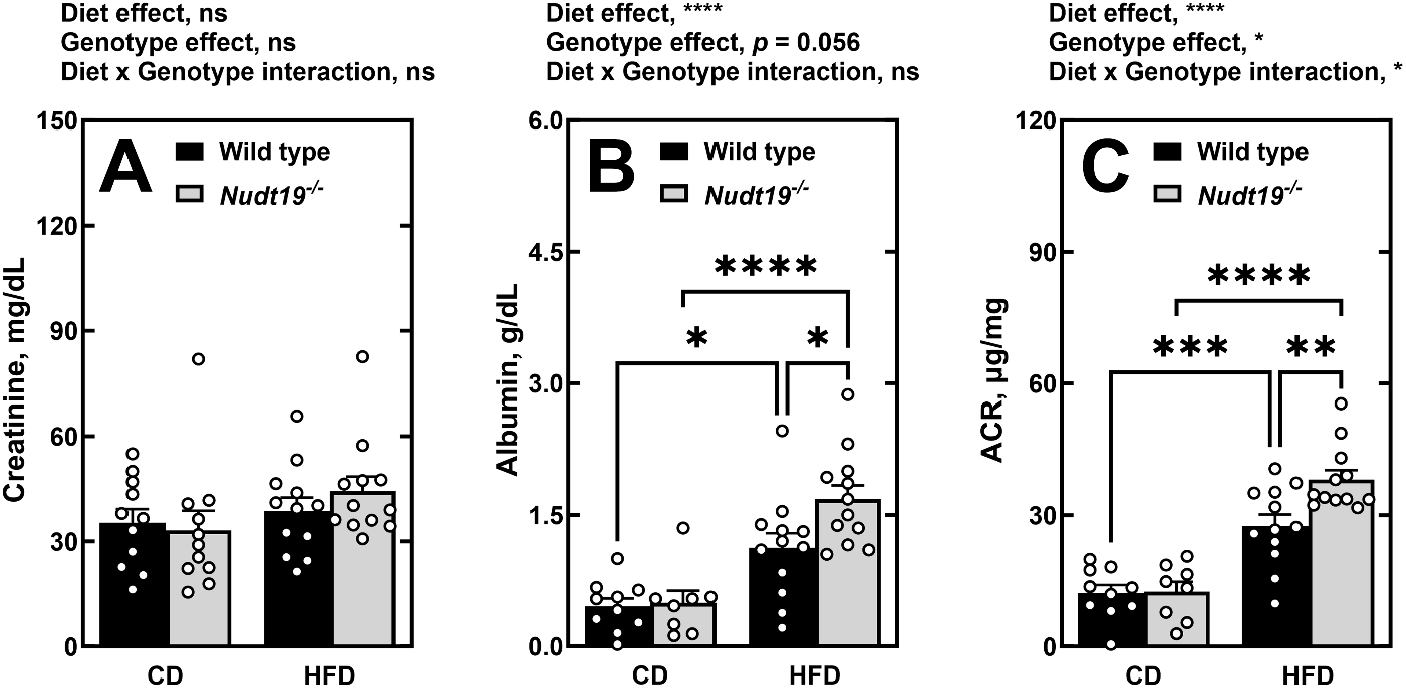
Deletion of *Nudt19* exacerbates the albuminuria induced by the HFD. (A) Creatinine, (B) albumin, and (C) albumin-to-creatinine ratio (ACR) in 24-hour urine samples collected from wild type and *Nudt19*^*-/-*^ mice fed the indicated diets. Data are shown as the mean (bars) of measurements from individual mice (circles) ± SEM. Two-way ANOVA; ns, not significant, *, p<0.05; **, p<0.01; ***, p<0.001; ****, p<0.0001.

### Effect of Nudt19 deletion on the kidney acyl-CoA pool composition, metabolome and proteome

NUDT19 hydrolyzes a variety of CoA species *in vitro*, and deletion of *Nudt19* in mice fed standard chow leads to a small but significant increase in total kidney CoA levels [28, 29]. These results are consistent with the peroxisomal CoASH/acyl-CoA pool being small compared to the amount of cofactor in the mitochondria and cytoplasm [40] and with the possibility that NUDT19 may only be able to access and hydrolyze the peroxisomal CoA pool. At the whole tissue levels, the kidneys of *Nudt19*^*-/-*^ mice fed the HFD showed similar total CoA levels (**Fig. 4A**) and acyl-CoA pool composition as wild type mice (**Fig. 4 B-D**). The only acyl-CoA that was significantly different between genotypes was isovaleryl/methylbutyryl/acetoacetyl-CoA, an intermediate in the metabolism of branched-chain amino acids or fatty acids, which was decreased in the *Nudt19*^*-/-*^ mice. Because local changes in peroxisomal acyl-CoA levels might be difficult to detect at the whole tissue level, we conducted global metabolomics and proteomics analyses to determine whether the absence of NUDT19 led to significant alterations in kidney metabolism.

**Figure 4.**
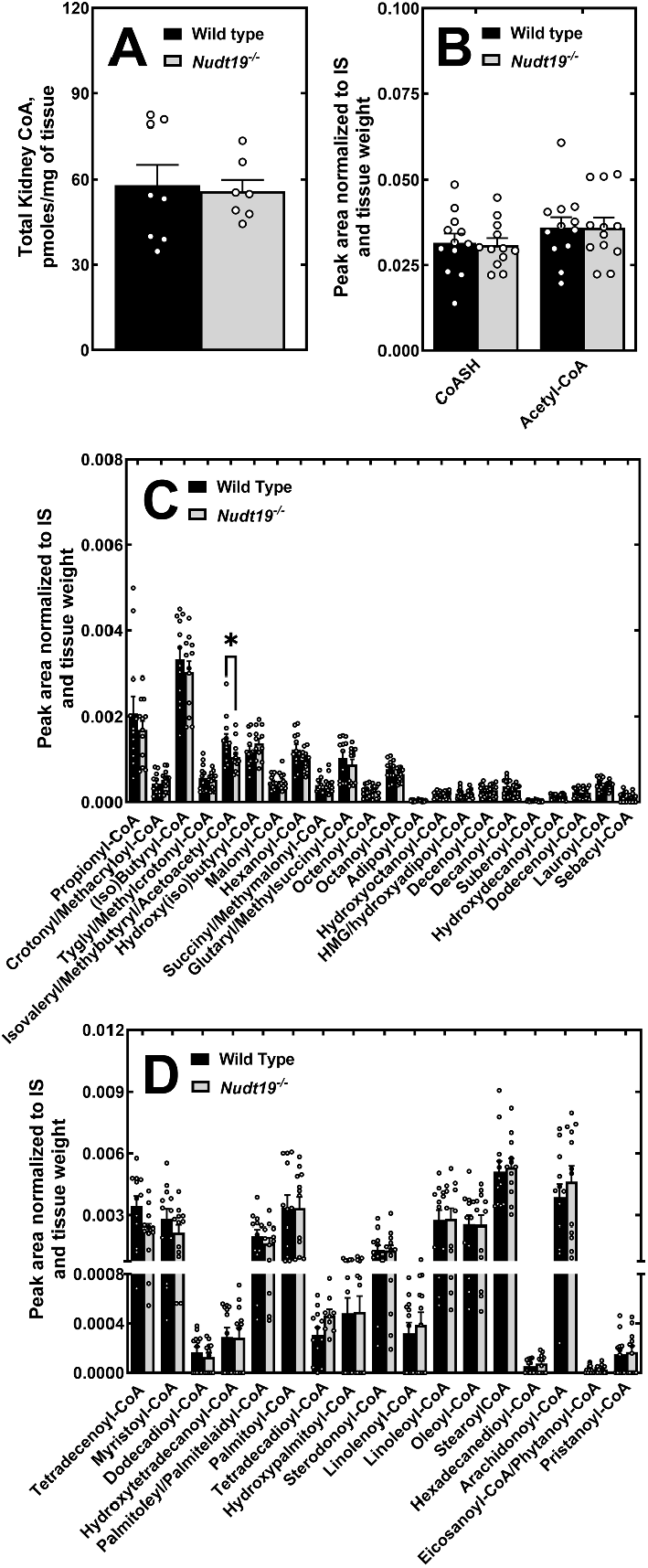
*Nudt19* deletion has no major effect on total kidney CoA levels and acyl-CoA composition. (A) Total CoA levels in the kidneys of mice fed the HFD. (B) CoASH and acetyl-CoA, (C) short- and medium-chain acyl-CoAs, and (D) long-chain acyl-CoAs. Data are shown as the mean (bars) of measurements from individual mice (circles) ± SEM. Student’s *t* test, * p<0.05.

In mice fed the HFD, untargeted metabolomics analysis identified 126 significantly changed metabolites out of the 839 known compounds detected (**Supplementary Table S1** and **Fig. 5**). Of the 126 compounds significantly changed, 55 were significantly increased and 71 were significantly decreased in the *Nudt19*^*-/-*^ mice (**Fig. 5A**). Pathway-enrichment analysis revealed that processes related to lipid metabolism, particularly non-esterified fatty acids (NEFA), were significantly altered in kidneys of mice lacking NUDT19 (**Fig. 5B**). Long-chain fatty acids exhibited a global decrease across all classes, including essential, monounsaturated, polyunsaturated, and branched-chain fatty acids (**Fig. 5C**), in correlation with reduced albumin reabsorption. Notably, the lower levels of branched-chain fatty acids may account for the decreased concentrations of short branched-chain acyl-CoA species, such as isovaleryl/methylbutyryl-CoA (**Fig. 4C**), which are generated in the peroxisomes through oxidation and shortening of branched-chain fatty acids. In addition to long-chain fatty acids, several monoacyl derivatives, including acylcholines, acyl ethanolamides, and monoacylglycerols, were also significantly reduced overall (**Fig. 5D**).

**Figure 5.**
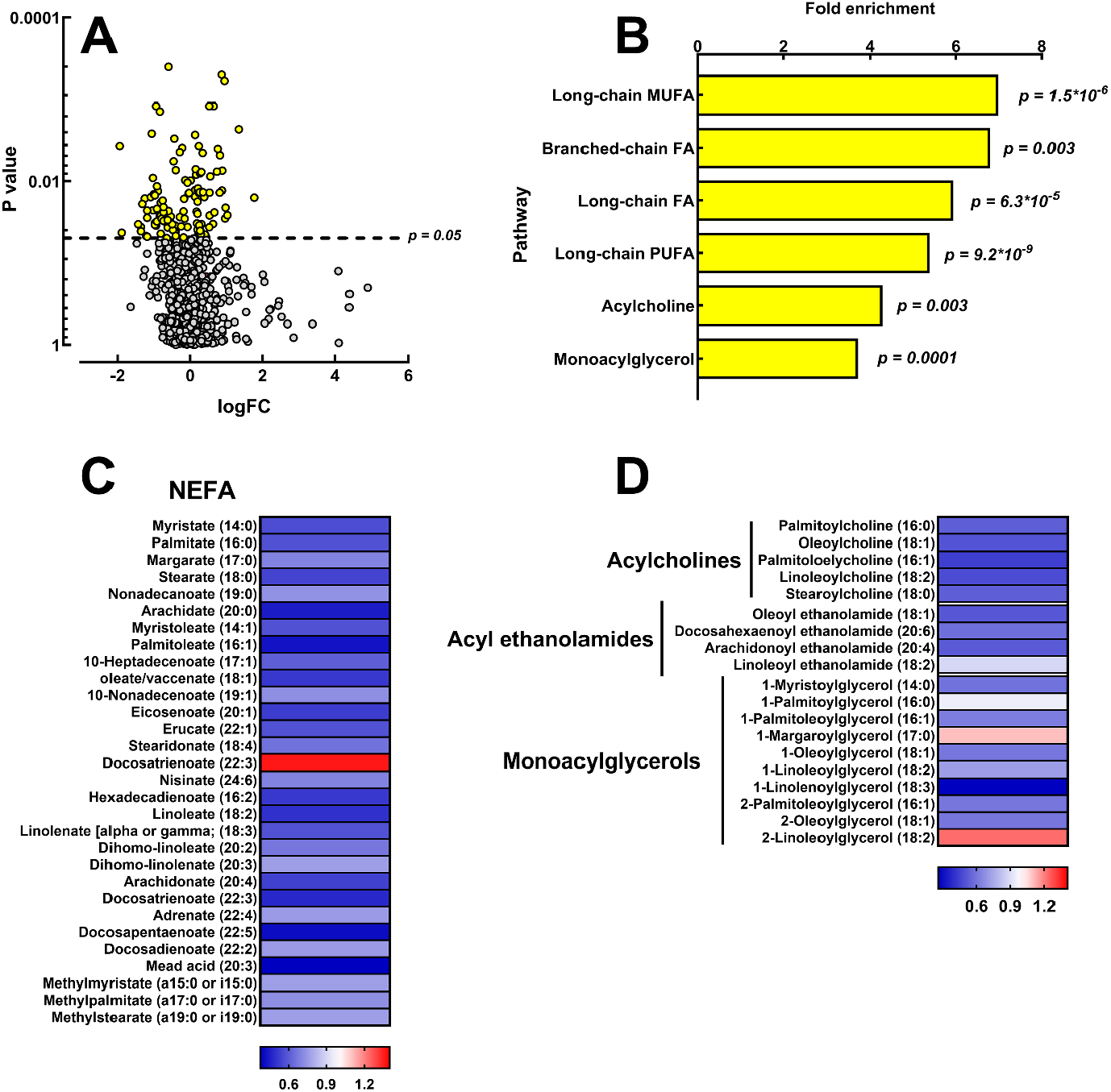
*Nudt19* deletion decreases kidney content of NEFA and monoacyl lipids. (A) Volcano plot of all metabolites detected by the untargeted metabolomics analysis, with significantly changed metabolites colored in yellow. (B) Pathway enrichment analysis, with p value derived from hypergeometric distribution test. (C-D) Heatmaps of all significantly changed (C) NEFA and (D) monoacyl lipids.

Global proteomics analysis of kidney cortex samples from mice fed the HFD identified 9 significantly altered proteins in the *Nudt19*^*-/-*^ mice, with 5 proteins increased and 4, including NUDT19, decreased. Although the small number of proteins precluded pathway enrichment analysis, 4 of the 5 elevated proteins were associated with lipid metabolism (**Fig 6A**). A fifth protein involved in lipid metabolism, the lipolysis-stimulated lipoprotein receptor (LSR, encoded by *Lsr*), was reduced by about 40% in the *Nudt19*^*-/-*^ kidneys (**Fig. 6A**). LSR binds low-density lipoproteins in the presence of NEFA and regulates the clearance of ApoB- and ApoE-containing lipoproteins from the blood [41, 42]. LSR localizes to the cell membrane, including tricellular tight junctions, where it regulates epithelial barrier function, and the nucleus [43, 44]. While LSR role in the kidneys is unclear, its deletion in proximal tubules is protective against unilateral ischemia-reperfusion injury [44].

**Figure 6.**
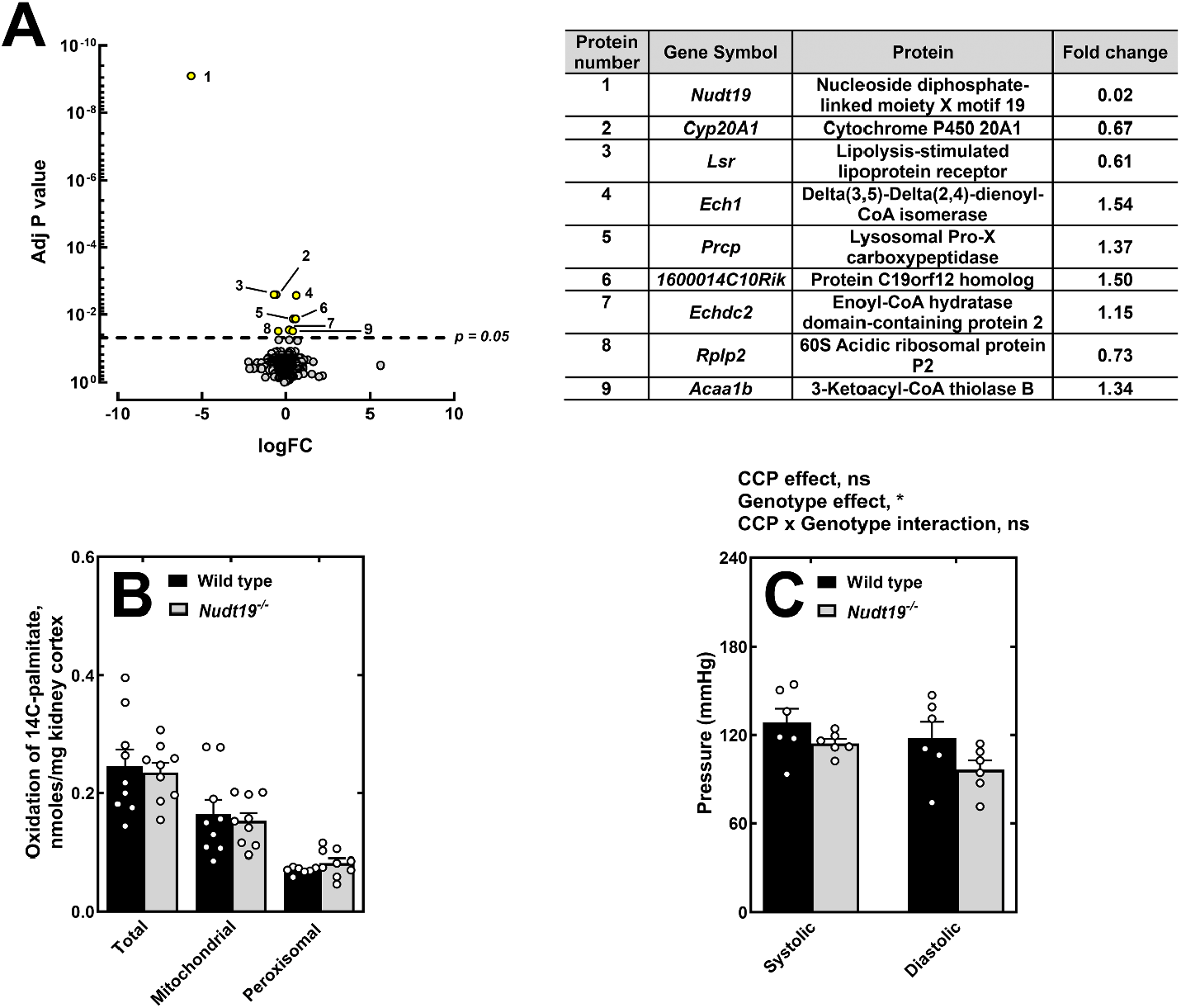
*Nudt19* deletion primarily affects the expression of proteins involved in lipid metabolism. (A) Volcano plot of proteins detected by tandem mass tag (TMT) proteomics, with significantly altered proteins highlighted in yellow and numbered. Corresponding protein identities and fold changes are listed in the accompanying table. (B) Measurement of ^14^C-palmitic acid oxidation in kidney slices. (C) Systolic and diastolic blood pressure in *Nudt19*^*-/-*^ and wild type mice fed a high-fat diet (HFD). Data in (B) and (C) are presented as the mean (bars) of individual measurements (circles) ± SEM. Student’s t test for (B) and two-way ANOVA for (C); *, p<0.05. CCP, cardiac cycle pressure.

Among the proteins increased in the *Nudt19*^*-/-*^ mice were Delta(3,5)-Delta(2,4)-dienoyl-CoA isomerase (ECH1, encoded by *Ech1*), which is involved in the oxidation of unsaturated fatty acids in both mitochondria and peroxisomes [45], and 3-ketoacyl-CoA thiolase B (THIKB, encoded by *Acaa1b*), an enzyme involved in the peroxisomal oxidation of straight-chain fatty acids [46]. Other elevated proteins included enoyl-CoA hydratase domain-containing protein 2 (ECHD2, encoded by *Echdc2*), a mitochondrial enzyme that shares limited sequence similarity with the β-oxidation enzyme enoyl-CoA hydratase [47], and protein C19orf12 homolog (encoded by *1600014CRik*). The latter is the mouse equivalent of the human protein C19orf12, which is mutated in patients with Mitochondrial membrane Protein-Associated Neurodegeneration (MPAN) [48]. Recent evidence indicate that this protein localizes to lipid droplets and regulates lipolysis and mitochondrial fatty acid oxidation in adipocytes [49]. To determine whether the increased expression of ECH1, THIKB, ECHD2 and protein C19orf12 homolog, combined with deletion of *Nudt19*, had a significant effect on fatty acid oxidation, we measure the rate of ^14^C-palmitic acid oxidation in kidney cortex slices, but no difference in mitochondrial or peroxisomal fatty acid oxidation was observed between genotypes (**Fig. 6B**).

The last enzyme whose levels were significantly increased in the kidneys of the *Nudt19*^*-/-*^ mice was lysosomal Pro-X carboxypeptidase (PRCP, encoded by *Prcp*). PRCP is a carboxypeptidase highly active at the acid pH usually found in collecting tubules and capable of cleaving angiotensin II, a peptide hormone known to interfere with albumin reabsorption, to angiotensin-(1-7) [50, 51]. PCRP-deficient mice show normal urinary sodium excretion but constitutively elevated blood pressure [52, 53]. Consistent with the increased levels of PCRP, measurement of systolic and diastolic blood pressure showed that *Nudt19*^*-/-*^ mice fed the HFD tended to have lower blood pressure values than wild type males, with a statistically significant genotype effect (**Fig. 6C**).

## Discussion

In this study, we examined the impact of a HFD on kidney metabolism in mice lacking NUDT19, a peroxisomal enzyme predominantly expressed in the kidneys. Our findings indicate that *Nudt19* deletion primarily affected lipid metabolism, leading to decreased levels of NEFA and select mono-acyl lipids. This phenotype was associated with a decrease in albumin reabsorption that, together with lower levels of LSR, could potentially explain the decrease in tissue NEFA. Disturbances in fatty acid metabolism in the *Nudt19*^*-/-*^ mice were independently supported by increased expression of ECH1, THIKB, ECHD2, enzymes involved in peroxisomal and mitochondrial β-oxidation, and C19orf12, which localizes to lipid droplets and regulates lipolysis [49]. In the context of *Nudt19* deletion, the increased expression of this protein, along with ECH1, THIKB, ECHD2, could represent a compensatory response to the reduction in endogenous NEFA. While ex vivo assays did not reveal significant changes in fatty acid oxidation rates between genotypes, these findings, together with recent evidence that *Nudt7* deletion alters bile acid and dicarboxylic fatty acid metabolism [26], underscore the importance of local acyl-CoA regulation for organ function.

The decrease in albumin reabsorption in *Nudt19*^*-/-*^ mice occurred despite lower NEFA levels, which are typically associated with lipotoxicity and kidney injury [54-56], and without detectable triglyceride or cholesterol accumulation at the whole-tissue level. While these findings do not rule out segment-or cell-type-specific lipid accumulation, they suggest that the relationship between lipid metabolism and kidney function may be more complex than previously thought, raising the possibility that lipid accumulation does not always precede albuminuria in the context of a HFD. Given the role of peroxisomes in lipid metabolism and their interactions with other organelles, it is possible that changes in peroxisomal function contribute to the observed effects on albumin reabsorption [57-61]. Indeed, while peroxisomes are not directly known to regulate albumin reabsorption, which occurs via receptor-mediated endocytosis, the *Nudt19*-dependent decrease in albumin reabsorption could be mediated by changes in lipid composition, sensing, or signaling in other subcellular compartments. Notably, this phenotype was accompanied by increased PRCP protein levels, which could represent a compensatory response to the higher ACR, as elevated PRCP activity would be expected to hydrolyze angiotensin II and counteract its inhibitory effect on albumin reabsorption [50, 51, 62]. Further studies are needed to clarify how peroxisomal acyl-CoA degradation affects albumin reabsorption under conditions of dietary fatty acid overload. Given that NUDT19 expression is reduced in kidney dysfunction or injury, including diabetic kidney disease [31] and ischemia-reperfusion [63], its loss may have broader implications for renal pathology. Analysis of published snRNA-seq datasets further supports this link, showing decreased *Nudt19* transcripts in the proximal tubules of mice subjected to unilateral ureteral obstruction (humphreyslab.com/SingleCell/) [64, 65]. Future studies could explore whether maintaining NUDT19 activity is sufficient to affect the kidney’s response to injury or diabetes. The relevance of NUDT19 may extend beyond kidney function, as emerging evidence links its expression to lipid metabolism and disease progression in multiple contexts. In prostate cancer, NUDT19 has been identified as part of a fatty acid metabolism-related gene signature with prognostic significance, where its increased expression correlates with disease progression [66]. Similarly, while NUDT19 is barely detectable in normal mouse liver [29], its expression is markedly upregulated in hepatocellular carcinoma (HCC), where it activates the mTORC1/P70S6K signaling pathway, promoting proliferation and cell migration [67]. Functionally, knocking down *Nudt19* in Hepa 1–6 cells enhances fatty acid oxidation and ATP production [68], further supporting its role in metabolic regulation. Collectively, these findings suggest that NUDT19 plays a broader role in cellular lipid metabolism and may contribute to disease pathogenesis in multiple organs.

## Abbreviations

(BSA): bovine serum albumin
(CD): low fat control diet
(HFD): high fat diet
(NEFA): non-esterified fatty acids
(TAG): triglycerides

## Acknowledgements

We thank Terence McManus from the WVU Metabolome Analysis Facility for his expert technical assistance.

## Funding and additional information

This work was supported by the WVU School of Medicine and the National Institutes of Health (NIH) (grant numbers R35GM119528 and R35GM119528-08S1), The Metabolome Core at West Virginia University was supported by the WV-CTSI program, which is funded by NIH grant 2U54GM104942. The content of this report is solely the responsibility of the authors and does not necessarily represent the official views of the National Institutes of Health.

## Supplementary Material

**Supplementary Figure S1.**
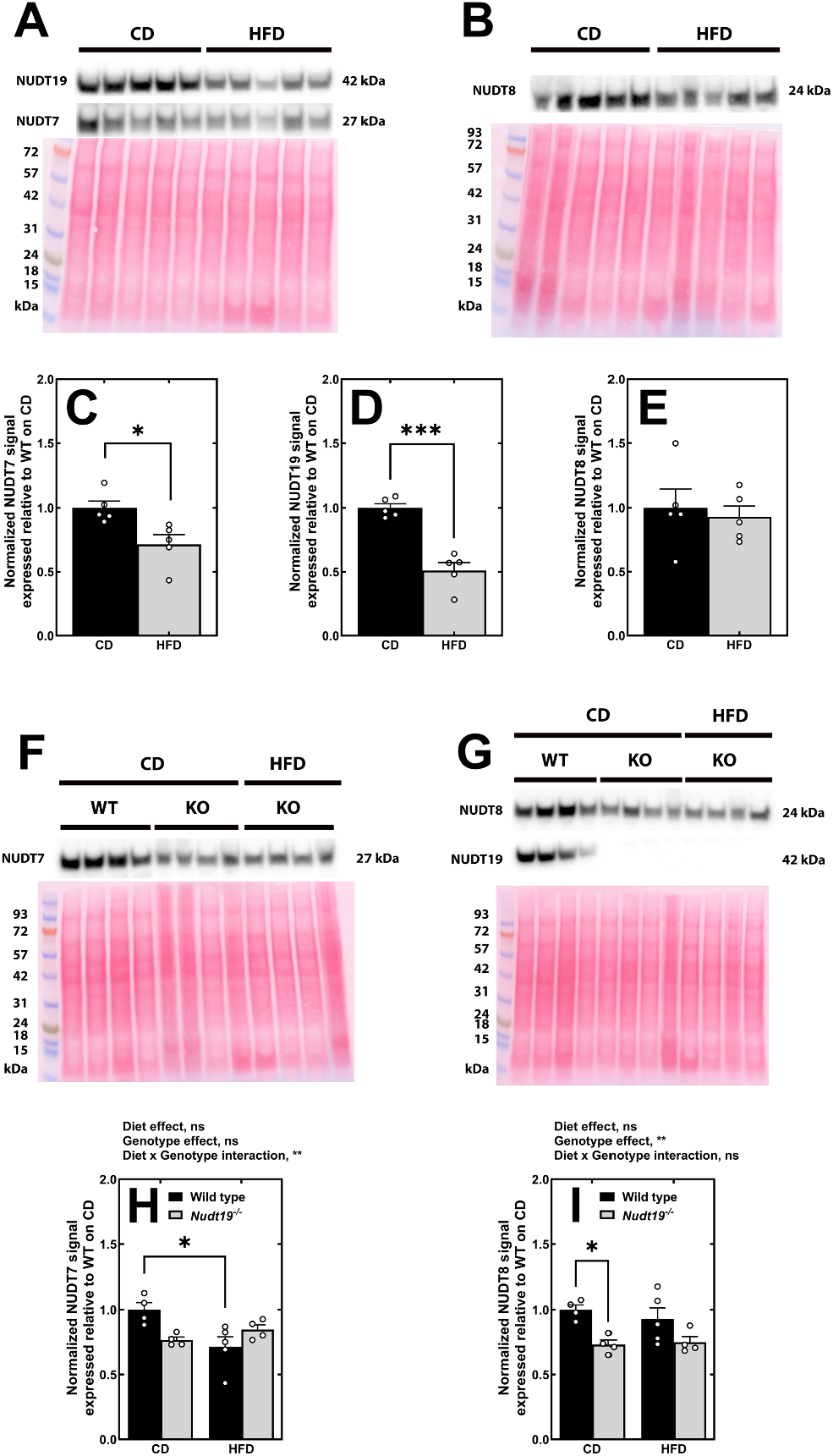
Effect of HFD and *Nudt19* deletion on the expression of NUDT7, NUDT8, and NUDT19 in the kidneys. (A) NUDT7 and NUDT19 protein levels in wild type (WT) mice fed the CD or the HFD. (B) NUDT8 protein levels in WT mice under the same conditions. (C–E) Quantification of Western blot signals. (F–G) Western blots detecting NUDT7, NUDT8, and NUDT19 in WT and Nudt19^−/−^ (KO) mice fed either the CD or HFD. (H–I) Corresponding quantification. Graphed data are shown as the mean (bars) of individual measurements (circles) ± SEM. Student’s *t* test for (C-D) and two-way ANOVA for (H, I); *, p<0.05; ***, p<0.01.

## References

1. Balaban RS, Mandel LJ. Metabolic substrate utilization by rabbit proximal tubule. An NADH fluorescence study. Am J Physiol. 1988;254(3 Pt 2):F407–16. doi: 10.1152/ajprenal.1988.254.3.F407. PubMed PMID: 3348418.

2. Elhamri M, Martin M, Ferrier B, Baverel G. Substrate uptake and utilization by the kidney of fed and starved rats in vivo. Ren Physiol Biochem. 1993;16(6):311–24. doi: 10.1159/000173777. PubMed PMID: 7506440.

3. Wirthensohn G, Guder WG. Renal substrate metabolism. Physiol Rev. 1986;66(2):469–97. doi: 10.1152/physrev.1986.66.2.469. PubMed PMID: 2938198.

4. Ross BD, Espinal J, Silva P. Glucose metabolism in renal tubular function. Kidney Int. 1986;29(1):54–67. doi: 10.1038/ki.1986.8. PubMed PMID: 3515015.

5. Lee JB, Peterhm. Effect of oxygen tension on glucose metabolism in rabbit kidney cortex and medulla. Am J Physiol. 1969;217(5):1464–71. doi: 10.1152/ajplegacy.1969.217.5.1464. PubMed PMID: 5346315.

6. Pfaller W. Structure function correlation on rat kidney. Quantitative correlation of structure and function in the normal and injured rat kidney. Adv Anat Embryol Cell Biol. 1982;70:1-106. PubMed PMID: 7058706.

7. Guder WG, Ross BD. Enzyme distribution along the nephron. Kidney Int. 1984;26(2):101–11. doi: 10.1038/ki.1984.143. PubMed PMID: 6094907.

8. Beard ME, Novikoff AB. Distribution of peroxisomes (microbodies) in the nephron of the rat: a cytochemical study. J Cell Biol. 1969;42(2):501–18. doi: 10.1083/jcb.42.2.501. PubMed PMID: 5792337; PubMed Central PMCID: PMCPMC2107674.

9. Rhodin JAG. Correlation of ultrastructural organization and function in normal and experimentally changed proximal convolutes tubule cells of mouse kidney [Doctoral]. Stockholm, Aktiebolaget Godvil: Karolinska Institutet; 1954.

10. Wanders RJ, Waterham HR, Ferdinandusse S. Metabolic Interplay between Peroxisomes and Other Subcellular Organelles Including Mitochondria and the Endoplasmic Reticulum. Front Cell Dev Biol. 2015;3:83. doi: 10.3389/fcell.2015.00083. PubMed PMID: 26858947; PubMed Central PMCID: PMCPMC4729952.

11. Lodhi IJ, Semenkovich CF. Peroxisomes: a nexus for lipid metabolism and cellular signaling. Cell Metab. 2014;19(3):380–92. doi: 10.1016/j.cmet.2014.01.002. PubMed PMID: 24508507; PubMed Central PMCID: PMC3951609.

12. Hunt MC, Tillander V, Alexson SE. Regulation of peroxisomal lipid metabolism: the role of acyl-CoA and coenzyme A metabolizing enzymes. Biochimie. 2014;98:45–55. doi: 10.1016/j.biochi.2013.12.018. PubMed PMID: 24389458.

13. Ding J, Loizides-Mangold U, Rando G, Zoete V, Michielin O, Reddy JK, et al. The peroxisomal enzyme L-PBE is required to prevent the dietary toxicity of medium-chain fatty acids. Cell Rep. 2013;5(1):248–58. doi: 10.1016/j.celrep.2013.08.032. PubMed PMID: 24075987.

14. Bergseth S, Poisson JP, Bremer J. Metabolism of dicarboxylic acids in rat hepatocytes. Biochim Biophys Acta. 1990;1042(2):182–7. doi: 10.1016/0005-2760(90)90005-i. PubMed PMID: 2302418.

15. Suzuki H, Yamada J, Watanabe T, Suga T. Compartmentation of dicarboxylic acid beta-oxidation in rat liver: importance of peroxisomes in the metabolism of dicarboxylic acids. Biochim Biophys Acta. 1989;990(1):25–30. doi: 10.1016/s0304-4165(89)80007-8. PubMed PMID: 2914148.

16. Leonardi R, Zhang YM, Rock CO, Jackowski S. Coenzyme A: back in action. Prog Lipid Res. 2005;44(2-3):125-53. doi: 10.1016/j.plipres.2005.04.001. PubMed PMID: 15893380.

17. Hirschey MD, Zhao Y. Metabolic Regulation by Lysine Malonylation, Succinylation, and Glutarylation. Mol Cell Proteomics. 2015;14(9):2308–15. doi: 10.1074/mcp.R114.046664. PubMed PMID: 25717114; PubMed Central PMCID: PMC4563717.

18. Resh MD. Fatty acylation of proteins: The long and the short of it. Prog Lipid Res. 2016;63:120–31. doi: 10.1016/j.plipres.2016.05.002. PubMed PMID: 27233110; PubMed Central PMCID: PMC4975971.

19. Sabari BR, Zhang D, Allis CD, Zhao Y. Metabolic regulation of gene expression through histone acylations. Nat Rev Mol Cell Biol. 2017;18(2):90–101. doi: 10.1038/nrm.2016.140. PubMed PMID: 27924077; PubMed Central PMCID: PMC5320945.

20. Gout I. Coenzyme A, protein CoAlation and redox regulation in mammalian cells. Biochem Soc Trans. 2018;46(3):721–8. doi: 10.1042/BST20170506. PubMed PMID: 29802218; PubMed Central PMCID: PMCPMC6008590.

21. Peng Y, Puglielli L. N-lysine acetylation in the lumen of the endoplasmic reticulum: A way to regulate autophagy and maintain protein homeostasis in the secretory pathway. Autophagy. 2016;12(6):1051–2. doi: 10.1080/15548627.2016.1164369. PubMed PMID: 27124586; PubMed Central PMCID: PMC4922439.

22. Trefely S, Huber K, Liu J, Noji M, Stransky S, Singh J, et al. Quantitative subcellular acyl-CoA analysis reveals distinct nuclear metabolism and isoleucine-dependent histone propionylation. Mol Cell. 2022;82(2):447–62 e6. doi: 10.1016/j.molcel.2021.11.006. PubMed PMID: 34856123.

23. Bulusu V, Tumanov S, Michalopoulou E, van den Broek NJ, MacKay G, Nixon C, et al. Acetate Recapturing by Nuclear Acetyl-CoA Synthetase 2 Prevents Loss of Histone Acetylation during Oxygen and Serum Limitation. Cell Rep. 2017;18(3):647–58. doi: 10.1016/j.celrep.2016.12.055. PubMed PMID: 28099844; PubMed Central PMCID: PMCPMC5276806.

24. Becuwe M, Bond LM, Pinto AFM, Boland S, Mejhert N, Elliott SD, et al. FIT2 is an acyl-coenzyme A diphosphatase crucial for endoplasmic reticulum homeostasis. J Cell Biol. 2020;219(10). doi: 10.1083/jcb.202006111. PubMed PMID: 32915949; PubMed Central PMCID: PMCPMC7659722.

25. Bond LM, Ibrahim A, Lai ZW, Walzem RL, Bronson RT, Ilkayeva OR, et al. Fitm2 is required for ER homeostasis and normal function of murine liver. J Biol Chem. 2023;299(3):103022. doi: 10.1016/j.jbc.2023.103022. PubMed PMID: 36805337; PubMed Central PMCID: PMCPMC10027564.

26. Vickers SD, Shumar SA, Saporito DC, Kunovac A, Hathaway QA, Mintmier B, et al. NUDT7 regulates total hepatic CoA levels and the composition of the intestinal bile acid pool in male mice fed a Western diet. JBiol Chem. 2023;299(1):102745. doi: 10.1016/j.jbc.2022.102745. PubMed PMID: 36436558; PubMed Central PMCID: PMCPMC9792899.

27. Shumar SA, Kerr EW, Fagone P, Infante AM, Leonardi R. Overexpression of Nudt7 decreases bile acid levels and peroxisomal fatty acid oxidation in the liver. J Lipid Res. 2019;60(5):1005–19. doi: 10.1194/jlr.M092676. PubMed PMID: 30846528; PubMed Central PMCID: PMCPMC6495166.

28. Ofman R, Speijer D, Leen R, Wanders RJ. Proteomic analysis of mouse kidney peroxisomes: identification of RP2p as a peroxisomal nudix hydrolase with acyl-CoA diphosphatase activity. Biochem J. 2006;393(Pt 2):537–43. doi: 10.1042/BJ20050893. PubMed PMID: 16185196; PubMed Central PMCID: PMC1360704.

29. Shumar SA, Kerr EW, Geldenhuys WJ, Montgomery GE, Fagone P, Thirawatananond P, et al. Nudt19 is a renal CoA diphosphohydrolase with biochemical and regulatory properties that are distinct from the hepatic Nudt7 isoform. J Biol Chem. 2018;293(11):4134–48. doi: 10.1074/jbc.RA117.001358. PubMed PMID: 29378847; PubMed Central PMCID: PMCPMC5857999.

30. Rheaume C, Barbour KW, Tseng-Crank J, Berger FG. Molecular genetics of androgen-inducible RP2 gene transcription in the mouse kidney. Mol Cell Biol. 1989;9(2):477-83. PubMed PMID: 2710112; PubMed Central PMCID: PMC362623.

31. Tserga A, Pouloudi D, Saulnier-Blache JS, Stroggilos R, Theochari I, Gakiopoulou H, et al. Proteomic Analysis of Mouse Kidney Tissue Associates Peroxisomal Dysfunction with Early Diabetic Kidney Disease. Biomedicines. 2022;10(2). Epub 20220120. doi: 10.3390/biomedicines10020216. PubMed PMID: 35203426; PubMed Central PMCID: PMCPMC8869654.

32. McLennan AG. The Nudix hydrolase superfamily. Cell Mol Life Sci. 2006;63(2):123–43. doi: 10.1007/s00018-005-5386-7. PubMed PMID: 16378245.

33. Kerr EW, Shumar SA, Leonardi R. Nudt8 is a novel CoA diphosphohydrolase that resides in the mitochondria. FEBS Lett. 2019;593(11):1133–43. doi: 10.1002/1873-3468.13392. PubMed PMID: 31004344; PubMed Central PMCID: PMCPMC6557688.

34. Ohkawa H, Ohishi N, Yagi K. Assay for lipid peroxides in animal tissues by thiobarbituric acid reaction. Anal Biochem. 1979;95(2):351–8. doi: 10.1016/0003-2697(79)90738-3. PubMed PMID: 36810.

35. Shumar SA, Fagone P, Alfonso-Pecchio A, Gray JT, Rehg JE, Jackowski S, Leonardi R. Induction of Neuron-Specific Degradation of Coenzyme A Models Pantothenate Kinase-Associated Neurodegeneration by Reducing Motor Coordination in Mice. PLoS One. 2015;10(6):e0130013. doi: 10.1371/journal.pone.0130013. PubMed PMID: 26052948; PubMed Central PMCID: PMC4460045.

36. Nesvizhskii AI, Keller A, Kolker E, Aebersold R. A statistical model for identifying proteins by tandem mass spectrometry. Anal Chem. 2003;75(17):4646–58. doi: 10.1021/ac0341261. PubMed PMID: 14632076.

37. Graw S, Tang J, Zafar MK, Byrd AK, Bolden C, Peterson EC, Byrum SD. proteiNorm - A User-Friendly Tool for Normalization and Analysis of TMT and Label-Free Protein Quantification. ACS Omega. 2020;5(40):25625–33. doi: 10.1021/acsomega.0c02564. PubMed PMID: 33073088; PubMed Central PMCID: PMCPMC7557219.

38. Huber W, von Heydebreck A, Sultmann H, Poustka A, Vingron M. Variance stabilization applied to microarray data calibration and to the quantification of differential expression. Bioinformatics. 2002;18 Suppl 1:S96–104. doi: 10.1093/bioinformatics/18.suppl_1.s96. PubMed PMID: 12169536.

39. Ritchie ME, Phipson B, Wu D, Hu Y, Law CW, Shi W, Smyth GK. limma powers differential expression analyses for RNA-sequencing and microarray studies. Nucleic Acids Res. 2015;43(7):e47. doi: 10.1093/nar/gkv007. PubMed PMID: 25605792; PubMed Central PMCID: PMCPMC4402510.

40. Horie S, Isobe M, Suga T. Changes in CoA pools in hepatic peroxisomes of the rat under various conditions. J Biochem. 1986;99(5):1345-52. PubMed PMID: 3711067.

41. Stenger C, Hanse M, Pratte D, Mbala ML, Akbar S, Koziel V, et al. Up-regulation of hepatic lipolysis stimulated lipoprotein receptor by leptin: a potential lever for controlling lipid clearance during the postprandial phase. FASEB J. 2010;24(11):4218–28. Epub 20100720. doi: 10.1096/fj.10-160440. PubMed PMID: 20647547.

42. Yen FT, Masson M, Clossais-Besnard N, Andre P, Grosset JM, Bougueleret L, et al. Molecular cloning of a lipolysis-stimulated remnant receptor expressed in the liver. J Biol Chem. 1999;274(19):13390–8. doi: 10.1074/jbc.274.19.13390. PubMed PMID: 10224102.

43. Masuda S, Oda Y, Sasaki H, Ikenouchi J, Higashi T, Akashi M, et al. LSR defines cell corners for tricellular tight junction formation in epithelial cells. J Cell Sci. 2011;124(Pt 4):548-55. Epub 20110118. doi: 10.1242/jcs.072058. PubMed PMID: 21245199.

44. Jiang M, Wang X, Chen Z, Wang X, An Y, Ding L, et al. Lipolysis-Stimulated Lipoprotein Receptor in Proximal Tubule, BMP-SMAD Signaling, and Kidney Disease. J Am Soc Nephrol. 2024;35(8):1016–33. Epub 20240529. doi: 10.1681/ASN.0000000000000382. PubMed PMID: 38809616; PubMed Central PMCID: PMCPMC11377808.

45. Zhang D, Liang X, He XY, Alipui OD, Yang SY, Schulz H. Delta 3,5,delta 2,4-dienoyl-CoA isomerase is a multifunctional isomerase. A structural and mechanistic study. J Biol Chem. 2001;276(17):13622–7. Epub 20010117. doi: 10.1074/jbc.M011315200. PubMed PMID: 11278886.

46. Nicolas-Frances V, Arnauld S, Kaminski J, Ver Loren van Themaat E, Clemencet MC, Chamouton J, et al. Disturbances in cholesterol, bile acid and glucose metabolism in peroxisomal 3-ketoacylCoA thiolase B deficient mice fed diets containing high or low fat contents. Biochimie. 2014;98:86–101. Epub 20131126. doi: 10.1016/j.biochi.2013.11.014. PubMed PMID: 24287293.

47. Du J, Li Z, Li QZ, Guan T, Yang Q, Xu H, et al. Enoyl coenzyme a hydratase domain-containing 2, a potential novel regulator of myocardial ischemia injury. J Am Heart Assoc. 2013;2(5):e000233. Epub 20131009. doi: 10.1161/JAHA.113.000233. PubMed PMID: 24108764; PubMed Central PMCID: PMCPMC3835224.

48. Gregory A, Klopstock T, Kmiec T, Hogarth P, Hayflick SJ. Mitochondrial Membrane Protein-Associated Neurodegeneration. In: Adam MP, Feldman J, Mirzaa GM, Pagon RA, Wallace SE, Amemiya A, editors. GeneReviews((R)). Seattle (WA) 1993.

49. Klingelhuber F, Frendo-Cumbo S, Omar-Hmeadi M, Massier L, Kakimoto P, Taylor AJ, et al. A spatiotemporal proteomic map of human adipogenesis. Nat Metab. 2024;6(5):861–79. Epub 20240402. doi: 10.1038/s42255-024-01025-8. PubMed PMID: 38565923; PubMed Central PMCID: PMCPMC11132986.

50. Caruso-Neves C, Kwon SH, Guggino WB. Albumin endocytosis in proximal tubule cells is modulated by angiotensin II through an AT2 receptor-mediated protein kinase B activation. Proc Natl Acad Sci U S A. 2005;102(48):17513–8. Epub 20051117. doi: 10.1073/pnas.0507255102. PubMed PMID: 16293694; PubMed Central PMCID: PMCPMC1297674.

51. Tojo A, Onozato ML, Kurihara H, Sakai T, Goto A, Fujita T. Angiotensin II blockade restores albumin reabsorption in the proximal tubules of diabetic rats. Hypertens Res. 2003;26(5):413–9. doi: 10.1291/hypres.26.413. PubMed PMID: 12887133.

52. Adams GN, LaRusch GA, Stavrou E, Zhou Y, Nieman MT, Jacobs GH, et al. Murine prolylcarboxypeptidase depletion induces vascular dysfunction with hypertension and faster arterial thrombosis. Blood. 2011;117(14):3929–37. Epub 20110204. doi: 10.1182/blood-2010-11-318527. PubMed PMID: 21297000; PubMed Central PMCID: PMCPMC3083303.

53. Maier C, Schadock I, Haber PK, Wysocki J, Ye M, Kanwar Y, et al. Prolylcarboxypeptidase deficiency is associated with increased blood pressure, glomerular lesions, and cardiac dysfunction independent of altered circulating and cardiac angiotensin II. J Mol Med (Berl). 2017;95(5):473–86. Epub 20170203. doi: 10.1007/s00109-017-1513-9. PubMed PMID: 28160049.

54. Cobbs A, Ballou K, Chen X, George J, Zhao X. Saturated fatty acids bound to albumin enhance osteopontin expression and cleavage in renal proximal tubular cells. Int J Physiol Pathophysiol Pharmacol. 2018;10(1):29–38. Epub 20180310. PubMed PMID: 29593848; PubMed Central PMCID: PMCPMC5871627.

55. Adeosun SO, Gordon DM, Weeks MF, Moore KH, Hall JE, Hinds TD, Jr., Stec DE. Loss of biliverdin reductase-A promotes lipid accumulation and lipotoxicity in mouse proximal tubule cells. Am J Physiol Renal Physiol. 2018;315(2):F323–F31. Epub 20180404. doi: 10.1152/ajprenal.00495.2017. PubMed PMID: 29631357; PubMed Central PMCID: PMCPMC6139518.

56. Castro BBA, Foresto-Neto O, Saraiva-Camara NO, Sanders-Pinheiro H. Renal lipotoxicity: Insights from experimental models. Clin Exp Pharmacol Physiol. 2021;48(12):1579–88. Epub 20210923. doi: 10.1111/1440-1681.13556. PubMed PMID: 34314523.

57. Schrader M, Kamoshita M, Islinger M. Organelle interplay-peroxisome interactions in health and disease. J Inherit Metab Dis. 2020;43(1):71–89. Epub 20190416. doi: 10.1002/jimd.12083. PubMed PMID: 30864148; PubMed Central PMCID: PMCPMC7041636.

58. Camoes F, Bonekamp NA, Delille HK, Schrader M. Organelle dynamics and dysfunction: A closer link between peroxisomes and mitochondria. J Inherit Metab Dis. 2009;32(2):163–80. Epub 20081212. doi: 10.1007/s10545-008-1018-3. PubMed PMID: 19067229.

59. Valm AM, Cohen S, Legant WR, Melunis J, Hershberg U, Wait E, et al. Applying systems-level spectral imaging and analysis to reveal the organelle interactome. Nature. 2017;546(7656):162–7. Epub 20170524. doi: 10.1038/nature22369. PubMed PMID: 28538724; PubMed Central PMCID: PMCPMC5536967.

60. Hua R, Cheng D, Coyaud E, Freeman S, Di Pietro E, Wang Y, et al. VAPs and ACBD5 tether peroxisomes to the ER for peroxisome maintenance and lipid homeostasis. J Cell Biol. 2017;216(2):367–77. Epub 20170120. doi: 10.1083/jcb.201608128. PubMed PMID: 28108526; PubMed Central PMCID: PMCPMC5294787.

61. Chu BB, Liao YC, Qi W, Xie C, Du X, Wang J, et al. Cholesterol transport through lysosome-peroxisome membrane contacts. Cell. 2015;161(2):291–306. doi: 10.1016/j.cell.2015.02.019. PubMed PMID: 25860611.

62. Afonso LG, Silva-Aguiar RP, Teixeira DE, Alves SAS, Schmaier AH, Pinheiro AAS, et al. The angiotensin II/type 1 angiotensin II receptor pathway is implicated in the dysfunction of albumin endocytosis in renal proximal tubule epithelial cells induced by high glucose levels. Biochim Biophys Acta Gen Subj. 2024;1868(10):130684. Epub 20240729. doi: 10.1016/j.bbagen.2024.130684. PubMed PMID: 39084330.

63. Gerhardt LMS, Liu J, Koppitch K, Cippa PE, McMahon AP. Single-nuclear transcriptomics reveals diversity of proximal tubule cell states in a dynamic response to acute kidney injury. Proc Natl Acad Sci U S A. 2021;118(27). doi: 10.1073/pnas.2026684118. PubMed PMID: 34183416; PubMed Central PMCID: PMCPMC8271768.

64. Wu H, Malone AF, Donnelly EL, Kirita Y, Uchimura K, Ramakrishnan SM, et al. Single-Cell Transcriptomics of a Human Kidney Allograft Biopsy Specimen Defines a Diverse Inflammatory Response. J Am Soc Nephrol. 2018;29(8):2069–80. Epub 20180706. doi: 10.1681/ASN.2018020125. PubMed PMID: 29980650; PubMed Central PMCID: PMCPMC6065085.

65. Wu H, Kirita Y, Donnelly EL, Humphreys BD. Advantages of Single-Nucleus over Single-Cell RNA Sequencing of Adult Kidney: Rare Cell Types and Novel Cell States Revealed in Fibrosis. J Am Soc Nephrol. 2019;30(1):23–32. Epub 20181203. doi: 10.1681/ASN.2018090912. PubMed PMID: 30510133; PubMed Central PMCID: PMCPMC6317600.

66. Zhao H, Wu T, Luo Z, Huang Q, Zhu S, Li C, et al. Construction and validation of a fatty acid metabolism-related gene signature for predicting prognosis and therapeutic response in patients with prostate cancer. PeerJ. 2023;11:e14854. Epub 20230206. doi: 10.7717/peerj.14854. PubMed PMID: 36778142; PubMed Central PMCID: PMCPMC9910187.

67. Lan C, Wang Y, Su X, Lu J, Ma S. LncRNA LINC00958 Activates mTORC1/P70S6K Signalling Pathway to Promote Epithelial-Mesenchymal Transition Process in the Hepatocellular Carcinoma. Cancer Invest. 2021;39(6-7):539-49. Epub 20210527. doi: 10.1080/07357907.2021.1929282. PubMed PMID: 33979257.

68. Gorigk S, Ouwens DM, Kuhn T, Altenhofen D, Binsch C, Damen M, et al. Nudix hydrolase NUDT19 regulates mitochondrial function and ATP production in murine hepatocytes. Biochim Biophys Acta Mol Cell Biol Lipids. 2022;1867(6):159153. Epub 20220331. doi: 10.1016/j.bbalip.2022.159153. PubMed PMID: 35367353.

